# Metamodal Coupling of Vibrotactile and Auditory Speech Processing Systems Through Matched Stimulus Representations

**DOI:** 10.1101/2021.05.04.442660

**Authors:** Srikanth R. Damera, Patrick S. Malone, Benson W. Stevens, Richard Klein, Silvio P. Eberhardt, Edward T. Auer, Lynne E. Bernstein, Maximilian Riesenhuber

## Abstract

It has been postulated that the brain is organized by “metamodal”, sensory-independent cortical modules implementing particular computations, leading to the intriguing hypothesis that brain areas can perform tasks (such as word recognition) not just in “standard” sensory modalities but also in novel sensory modalities. Yet, evidence for this theory, especially in neurotypical subjects, has been variable. We hypothesized that effective metamodal engagement of a brain area requires congruence between the novel and standard sensory modalities not only at the task level (e.g., “word recognition”) but critically also a match at the algorithmic level (in Marr’s terminology), i.e., at the level of neural representation of the information of interest. To test this hypothesis, we trained participants to recognize vibrotactile versions of auditory words using two encoding schemes. The vocoded approach preserved the dynamics and representational similarities of auditory speech while the token-based approach used an abstract phoneme-based code. Although both groups learned the vibrotactile word recognition task, only in the vocoded group did trained vibrotactile stimuli recruit the auditory speech network and lead to increased coupling between somatosensory and auditory speech areas. In contrast, the token-based encoding appeared to rely on paired-associate learning. Thus, matching neural input representations is a critical factor for assessing and leveraging the metamodal potential of cortical modules.

## Introduction

The dominant view of brain organization revolves around cortical areas dedicated for processing information from specific sensory modalities. However, emerging evidence over the past two decades has led to the idea that many sensory cortical areas are “metamodal”, i.e., characterized by computations that are task-specific, yet invariant to sensory modality (Heimler et al., 2015; Pascual-Leone and Hamilton, 2001). Early evidence for sensory modality-invariant processing in cortical areas comes from cross-modal plasticity studies in sensory-deprived populations (Rauschecker, 1995; Rauschecker et al., 1992; Sadato et al., 1996; Théoret et al., 2004). These studies showed that cortical areas traditionally considered to be dedicated to unisensory processing could be recruited by stimuli from another, non-preferred sensory modality. Interestingly, although broad swaths of cortex no longer received the normal sensory input in these populations, non-preferred modality stimuli activated the specific cortical areas normally relevant for a particular task in the preferred sensory modality, such as localization and recognition (Amedi et al., 2007; Bi et al., 2016; Bola et al., 2017, 2020; Lomber et al., 2010; Meredith et al., 2011; Ptito et al., 2005; Reich et al., 2011; Renier et al., 2014; Striem-Amit et al., 2012). This task-specific cross-modal engagement is thought to reflect a functional unmasking of existing anatomical connections. Importantly, there is evidence for cross-modal engagement of traditionally unisensory areas even in neurotypical individuals (Amedi et al., 2007; Renier et al., 2005, 2010; Siuda-Krzywicka et al., 2016) – thereby opening the door to recruiting previously established sensory processing pathways for novel sensory modalities. A prime example of this process is reading, which is initially thought to recruit auditory speech processing pathways through grapheme-to-phoneme conversion (Pugh et al., 2001), and the same idea has given rise to promising therapeutic applications such as sensory substitution devices (SSDs, which, for instance enable processing of visual information in blind individuals by translating camera input to acoustic stimuli (Bach-y-Rita and Kercel, 2003; Meijer, 1992). Yet, other studies (Benetti et al., 2017, 2020; Bola et al., 2017; Fairhall et al., 2017; Mattioni et al., 2020; Pietrini et al., 2004; Twomey et al., 2017; Vetter et al., 2020) have failed to find or have found far less robust evidence of cross-modal engagement in neurotypical subjects, raising the critical question of the conditions under which a particular sensory area can be successfully recruited for “metamodal” processing.

Current theories for the “metamodal” brain differ on the necessary requirements for the engagement of a sensory area. Prior studies (Amedi et al., 2002, 2007; Striem-Amit et al., 2012) have emphasized that metamodal engagement of a cortical area, in addition to the presence of task-relevant connectivity (Hannagan et al., 2015; Mahon and Caramazza, 2011; Saygin et al., 2012, 2016), depends on a correspondence of preferred and non-preferred modality stimuli. Yet, it has been challenging to elucidate the nature of this necessary “correspondence” for metamodal engagement. For example, the fact that (visual) shape-selective ventral occipito-temporal cortex (VOTC) is engaged in subjects trained on a visual-to-auditory SSD designed to transform visual images to sounds has been used to argue that this region responds to any modality that conveys shape information (Amedi et al., 2007; Hannagan et al., 2015; Heimler et al., 2015; Striem-Amit et al., 2012). Consequentially, failures to find metamodal engagement are due to an absence of shape information in the stimuli encoded in the non-standard modality. Yet, in the absence of a clear definition of auditory “shape” such theories face reverse inference challenges and are difficult to test, let alone falsify. In terms of Marr’s levels of analysis (Computational, Algorithmic, Implementational), what is needed is a consideration of the Algorithmic level that serves as a crucial bridge between the Computational level of task goals, and the Implementational level of neural responses (Barsalou, 2017; Hagoort, 2020). Here we propose that successful metamodal coupling requires not just correspondence at the Computational/task level, but also a match of the novel modality representation to the standard modality representation at the Algorithmic level. This theory concretizes and makes testable the helpful yet vague notion of “correspondence” between preferred and non-preferred modality representations by tying it to a correspondence of representational spaces. A failure to achieve this *representational* correspondence would then be predicted to impede or even preclude metamodal engagement.

In the present study, we test the hypothesis that cross-sensory recruitment of existing learned sensory processing pathways critically depends on this representational match. Specifically, we focused on auditory-to-tactile sensory substitution. This field has a long history dating back to the invention of the Tadoma method (Alcorn, 1945) – a method whereby deaf individuals learn to perceive auditory speech received via vibrotactile (VT) input from their fingers which are placed over the articulators of a speaker. Over a century of work on auditory-to-tactile sensory substitution has led to the development of VT speech aids (Gault, 1924, 1926). These devices have been used successfully to teach both deaf and hearing individuals to recognize auditory speech through touch (Bernstein et al., 1991; Brooks and Frost, 1983; Cieśla et al., 2019). Here we used such a device to train two neurotypical groups of adult subjects on the same word recognition task, with each group being trained with one of two auditory-to-VT sensory substitution algorithms. One algorithm was designed to preserve as much of the temporal dynamics of auditory speech as possible (“vocoded speech”), aiming to achieve a neural congruence between vibrotactile speech stimuli and auditory speech representations in brain areas that are part of the auditory speech system. The other algorithm (“token”) used a code in which specific VT patterns corresponded to specific phonetic features (Chomsky and Halle, 1968; Reed et al., 2018). Interestingly, at the behavioral level, subjects in both algorithm groups learned to associate VT patterns with spoken words at an equivalent level after training. However, fMRI analyses revealed critical differences in the cross-modal recruitment of brain areas between the two groups, with only the vocoded encoding group showing metamodal engagement of auditory speech processing areas, specifically the areas whose neural representations of auditory speech representation well matched the representational similarity of the vibrotactile word stimuli. Consistent with these findings, functional connectivity analyses showed that increased coupling between the auditory and somatosensory cortex after training also depended on the nature of the input representations produced by the different VT algorithms. These findings suggest that metamodal engagement of a cortical area is dependent not only on its task-relevant anatomical connectivity but also on the match at the level of representational encoding between the standard and novel modalities. Adopting the nomenclature of David Marr’s levels (Marr, 1982), our data show that a mere congruence at the highest, computational level (e.g., VT stimuli corresponding to auditory words) is insufficient for metamodal engagement. Rather, metamodal coupling requires a congruence at the algorithmic level (e.g., a match in neural representations). Thus, our study not only critically advances our understanding of metamodal engagement and thus general principles of brain organization, but also opens the door to designing more efficient sensory substitution algorithms that better interface with existing cortical processing pathways (as in the present study, where the algorithmically matched vocoded speech representation conveyed ∼1.2 times as much information per unit time than the non-matched one).

## Materials and Methods

### Participants

We recruited a total of 22, right-handed, healthy, native English speakers in this study (ages 18-27, 12 females). Georgetown University’s Institutional Review Board approved all experimental procedures, and written informed consent was obtained from all subjects before the experiment. We excluded 4 subjects from the auditory scan due to excessive motion (>20% of volumes) and 2 subjects from the vibrotactile (VT) scans because they failed to complete the training. As a result, we analyzed a total of 18 subjects for the auditory scans and 20 for the VT scans.

### Stimulus Selection

A set of word stimuli was developed according to the following criteria: 1) short monosyllabic stimuli (∼4 phonemes); 2) only contain phonemes from a limited subset of English consonants (8 consonants and 6 vowels); 3) set containing items predicted to be perceptually unique and therefore learnable; and 4) words that span the VT vocoder perceptual space (see below). To develop the set meeting these criteria we utilized a computational modeling approach based on the methods described in (Auer and Bernstein, 1997). Existing tactile consonant and vowel perceptual identification data (Bernstein, unpublished) were used in combination with the PhLex lexical database (Seitz, Bernstein, Auer, & MacEachern, 1998) to model the lexical perceptual space. In outline, the modeling steps are: (1) Transform phoneme identification data into groupings of phonemes as a function of a set level of dissimilarity; (2) Re-transcribe a phonemically transcribed lexical database so that all of the words are represented in terms of only the phonemic distinctions across groupings; and (3) Collect words that are identical under the re-transcription and count how many are in each collection. In this study, the lexical equivalence class (LEC) size –the number of words in a collection—was set to three. Only words that were accompanied by three or fewer other words following re-transcription were considered candidates for the study. Words in smaller LECs are predicted to be perceptually easier (more unique) than words in larger LECs, which offer more opportunities for confusions.

The set of words meeting the first three criteria was further examined as a function of consonants and vowel patterns to identify the largest pool of potential stimulus words. Three consonant (C) and vowel (V) segment patterns (CVC, CCVC, and CVCC) were selected for the final stimulus set. The words with these segment patterns were then examined in relation to the predicted VT vocoder perceptual space. The tactile identification confusion matrices were transformed into phoneme distance matrices using a phi-square transform (Iverson et al., 1998). Within a segment pattern, all word-to-word distances were computed as the sum of the pairwise phoneme distances. The word distance matrix was then submitted to multidimensional scaling to facilitate two-dimensional visualization of the lexical space. Close pairs were selected with goal of achieving distributed coverage in each of the three lexical spaces (CVC, CVCC, and CCVC). For each close pair, a third more distant word was chosen that provided a bridge to other pairs in the space. Final selection was based on the word-to-word computed distances using phi-square distances rather than the multidimensional space as clear warping was present due to the reduction of dimensionality.

This resulted in 60 total words or 20 sets of triples. We trained subjects to associate 30 words (10 triplets) with their corresponding VT tokens. In the RSA scans we used 15 (5 triplets) of these trained words of which 9 belonged to the CVCC, 3 to the CCVC and 3 to the CVC lexical classes (Fig. 1B).

**Figure 1:**
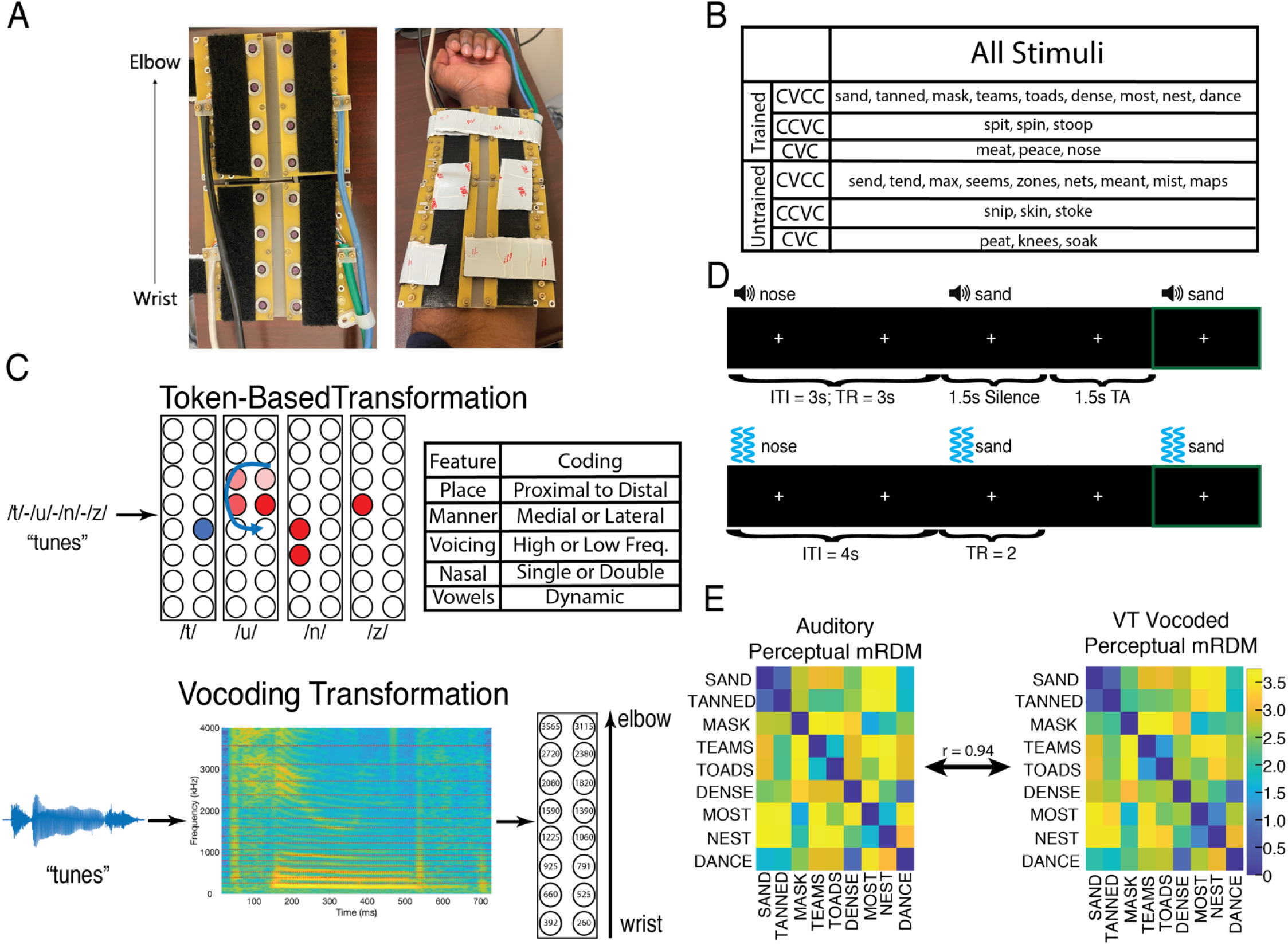
VT hardware, speech-to-tactile transformation algorithms, stimuli, fMRI experimental design, and model dissimilarity matrix. (A) Fourteen-channel MRI-compatible VT stimulator. (B) Shows the breakdown of the 30 words used in the study. The auditory scan used all the words, and subjects were trained on half of the words (“trained” set). Words were further broken down by their syllable structure (9 CVCC, 3 CCVC, and 3 CVC words). (C) Shows the two transformations used to convert spoken words into tactile stimulation patterns. The token-based approach (top) assigns each phoneme a distinct VT pattern (see Methods section for more details). The vocoding approach (bottom) focuses on preserving the temporal dynamics between the auditory and VT stimuli. (D) Shows the auditory (top) and VT (bottom) fMRI one-back paradigms used in the study. In both paradigms, subjects focused on a central fixation cross, and pressed a button in their left hand if they heard or felt the same stimulus twice in a row. (E) The auditory and VT vocoded perceptual model representational dissimilarity matrix (mRDM) for the 9 CVCC trained words. The high correlation (r = 0.94) between mRDMs provide evidence for the targeted close representational congruence between auditory and VT vocoded stimuli.

### Behavioral Training

The training paradigm used an N-alternative forced choice (N-AFC) task and a leveling system organized in sets of 3 to facilitate training progression. In each set of 3 levels, the number of choices (N) in the N-AFC task was kept constant, but the choices themselves were increasingly confusable. The number of choices N was increased by 1 when progressing between each set of 3 levels. The first level utilized a 2-AFC task, and the final level (level 15) utilized an 8-AFC task. An accuracy of 80% was required to advance to the next level. Subjects performed each training session in a quiet room while listening to an auditory white noise stimulus through over-the-ear headphones. Auditory white noise was presented in order to mask the mechanical sound of the VT stimulation. At the beginning of each trial, the orthographic labels for the word choices were displayed on the screen, and a VT stimulus was played after a short delay. Participants then indicated which label corresponded to the VT stimulus. Feedback was given after each trial, as well as an opportunity to replay any of the word choices. Subjects completed a total of 6 training sessions, followed by a post-training fMRI scan. After their post-training fMRI scan, subjects performed a final 10-AFC task.

### Description of VT Device

A (20cm x 11.0 cm) 16-channel MRI-compatible vibrotactile stimulator array was organized as 2 rows of 8 stimulators (Fig. 1A), with center-to-center stimulator spacing of 2.54 cm. To ensure that the stimulators would maintain contact with the volar forearm, the array comprised four rigid modules connected with stiff plastic springs. Velcro straps were used to mount the device to the arm firmly while bending the array to conform to the arm’s shape. With no applied voltage to the piezoelectric bimorphs, the contactors were flush with the circuit board surface facing the skin. During operation, a constant +57-V voltage applied to all stimulators retracted the contactors into the surround, and each applied -85-V pulse drove the contactor into the skin. All pulses were identical. The drive signal was a square wave, with a pulse time of 2 ms, and with unpowered intervals of 1ms between power reversals to protect the switching circuitry. The display’s control system comprised the power supplies (-85V, +57V), high voltage switching circuits to apply these voltages to the piezoelectric bimorphs, and a digital control system that accepted from a controlling computer’s serial COM port the digital records specifying a stimulus (comprising the times and channels to output pulses on), and a command to initiate stimulus output.

### VT Vocoded Speech Encoding

This real-time vocoder was used to convert acoustic speech signals into VT stimuli. The initial stage of the vocoder comprised a bank of filters whose output power was used to control the output of VT pulses. The VT display (Fig. 1A) used a frequency-to-place mapping algorithm: The energy passed by each filter of the vocoder was used to modulate the vibration of a specific MRI-compatible transducer on the 16-channel VT device (Fig. 1A and 1C) placed on the volar forearm (Malone et al., 2019). Low frequencies mapped to transducers near the wrist, and higher frequencies mapped to transducers near the elbow. If the energy within a given filter exceeded a fixed threshold at a given time point, a VT pulse was emitted from the corresponding transducer. The basic hardware design and software algorithms for the vocoder are referred to in (Bernstein et al., 1991) as the “GULin” vocoder algorithm. Briefly, 16 bandpass filters with frequencies centered at 260, 392, 525, 660, 791, 925, 1060, 1225, 1390, 1590, 1820, 2080, 2380, 2720, and 3115 Hz, with respective bandwidths of 115, 130, 130, 130, 130, 130, 145, 165,190, 220, 250, 290, 330, 375, and 435 Hz. An additional high-pass filter with cutoff 3565 Hz is also used. The energy detected in each band is used to amplitude-modulate a fixed-frequency sinewave at the center frequency of that band (and at 3565□Hz in the case of the high-pass filter). The combination of the 16 sinewaves comprises the vocoded acoustic signal, and the resulting activation pattern over the 16 transducers constituted its vibrotactile instantiation.

### Token-based VT Speech Encoding

The same 16-channel VT device was used to present subjects with the token-based stimuli. Token-based stimuli were constructed based on prior work (Reed et al., 2018) and reflect the idea that spoken words can be described as a string of phonemes. Phonemes in turn can be uniquely described by a set of phonetic features. Therefore, each phonetic feature was assigned a unique VT pattern. In this study, we used place, manner, and voicing features to describe phonemes (Fig. 1C). Place was coded as patterns that occurred either proximal or distal to the wrist. Stop and fricative manner features were codded as patterns that occurred either medial or lateral to the body respectively. The nasal manner feature was distinguished by driving two channels instead of one for stops and fricatives. Voicing was coded as either driving high frequency vibrations (250Hz) or low frequency vibrations (100Hz). Vowels were coded in a similar feature-based manner, but were dynamic stimuli (e.g. swirls and sweeps) whereas consonants were static. Importantly, all consonant patterns lasted 120ms and all vowel stimuli lasted 220ms and there was a 100ms gap between each pattern. As a result, token-based stimuli were either 660ms or 880ms long. CVCC trained token-based stimuli used in fMRI analyses were 880ms long while their VT vocoded counterparts had a mean duration of 727ms and standard deviation of 91.6ms. A paired t-test revealed that token-based stimuli were significantly longer (*t*(8) = 4.99; p = 0.001) than their vocoded counterparts. Thus, not only did VT vocoded but not token based stimuli preserve the temporal dynamics found in auditory speech, but they also conveyed more information per unit time.

### Auditory Scan

#### fMRI Experimental Procedures

EPI images from nine event-related runs were collected using a clustered acquisition paradigm. Within each run, 30 words were presented three times in random order for a total of 90 trials. Each trial was 3s long and started with 1.5s of volume acquisition followed by the auditory word (during the silent period, see below, “Data Acquisition”; Fig. 1D). To maintain attention, subjects performed a 1-back task in the scanner: Subjects were asked to press a button in their left hand whenever the same word was presented on two consecutive trials. These catch trials comprised ten percent of the trials in each run. Furthermore, an additional ten percent of trials were null trials. During these trials, which lasted for 3s, no words were presented. In total, there were 118 trials per run, with each trial lasting 3s for a total of 354s, plus an additional 15s fixation at the start of the run. Thus, in total each run lasted 369s and the session lasted 43min.

#### Data Acquisition

MRI data were acquired at the Center for Functional and Molecular Imaging at Georgetown University on a 3.0 Tesla Siemens Trio Scanner. We used whole-head echo-planar imaging sequences (flip angle = 90°, TE = 30 ms, FOV = 205, 64x64 matrix) with a 12-channel head coil. A clustered acquisition paradigm (TR = 3000 ms, TA = 1500 ms) was used such that each image was followed by an equal duration of silence before the next image was acquired. 28 descending axial slices were acquired in descending order (thickness = 3.5 mm, 0.5 mm gap; in-plane resolution = 3.0x3.0 mm^2^). This sequence was used in previous auditory studies from our lab (Chevillet et al., 2013). A T1-weighted MPRAGE image (resolution 1x1x1mm^3^) was also acquired for each subject.

### VT Scan

#### fMRI Experimental Procedures

EPI images from six event-related runs were collected. Within each run 30 stimuli (15 from the training set and 15 additional words) were presented three times in random order for a total of 90 trials. A 4 second intertrial interval was used (Fig. 1D). As in the auditory scan, to maintain attention, subjects performed a 1-back task in the scanner: Subjects were asked to press a button in their left hand whenever the same stimulus was presented on two consecutive trials. These catch trials comprised ten percent of the trials in each run. Furthermore, an additional ten percent of trials were null trials during which subjects were presented with a blank screen for 3s. In total, there were 111 trials per run with each trial lasting 4s for a total of 444s plus an additional 10s fixation at the start and end of the run. Thus, in total each run lasted 464s and the session lasted 46min.

#### Data Acquisition

MRI data were acquired at the Center for Functional and Molecular Imaging at Georgetown University on a 3.0 Tesla Siemens Trio Scanner. We used whole-head echo-planar imaging sequences (TR = 2000ms, flip angle = 90°, TE = 30 ms, FOV = 205, 64x64 matrix) with a 12-channel head coil. 33 interleaved descending axial slices were acquired (thickness = 3.5 mm, 0.5 mm gap; in-plane resolution = 3.0x3.0 mm^2^). A T1-weighted MPRAGE image (resolution 1x1x1mm^3^) was also acquired for each subject.

#### fMRI Data Preprocessing

Image preprocessing was performed in SPM12 (http://www.fil.ion.ucl.ac.uk/spm/software/spm12/) and AFNI. The first four acquisitions of each run were discarded to allow for T1 stabilization, and the remaining EPI images were slice-time corrected to the middle slice for the VT scans. No slice-time correction was performed for the auditory scans due to using a clustered acquisition paradigm due to temporal discontinuities between successive volumes (Perrachione and Ghosh, 2013). These images were then spatially realigned and submitted to the AFNI *align_epi_anat.py* function to co-register the anatomical EPI images for each subject. This was used because, upon inspection, it provided better registration between the anatomical and functional scans than the corresponding SPM12 routine.

### Anatomical Preprocessing

Freesurfer (Fischl et al., 1999) was used to reconstruct cortical surface models including an outer pial and inner white-matter surface. These surfaces were then brought into the SUMA environment and fit to a standardized meshe based on an icosahedron with 64 linear divisions using AFNI’s MapIcosehedron command (Oosterhof et al., 2011; Saad and Reynolds, 2011). This procedure yielded 81,924 nodes for each participant’s whole-brain cortical surface mesh. Each node on the standard mesh corresponds to the same location across subjects – thereby allowing node-wise group-level analysis. This improved the spatial resolution of our analyses since interpolation of the functional data is unnecessary (Oosterhof et al., 2011).

### Representational Similarity Analysis (RSA)

#### Constructing Model Representational Dissimilarity Matrices (mRDMs)

Two candidate mRDMs were generated: an auditory perceptual mRDM, and a VT vocoded perceptual mRDM. These mrDMs were generated by modifying an edit mRDM which was generated using an edit distance metric between word pairs in the stimulus set. Here, 1 edit was considered a substitution, insertion, or deletion of a single phoneme. Edit distances are frequently used with highly intelligible speech, for which there are no phoneme-to-phoneme dissimilarity data, and when more refined segment-to-segment distances are not available as was the case for the VT token-based algorithm. Furthermore, recent work (Kell et al., 2018) has shown that the representational format captured by the edit distance matches those found in both higher order STG speech regions and speech recognition-specific representations learned in later layers of a deep neural network. The auditory and VT vocoded perceptual mRDMs were similarly created using an edit distance but now weighting phoneme edit by either its auditory or VT vocoded perceptual confusability. Auditory and VT vocoded perceptual phoneme confusability was derived from a behaviorally measured perceptual auditory and VT vocoded phoneme identification task. This confusability was transformed into a distance measure using a phi-square transform (Iverson et al., 1998). Word-to-word distances were computed as the sum of the pairwise phoneme distances for all the position-specific phoneme pairs in each of the possible pairs of stimulus words. Given the difficulty of estimating a distance swap between consonants and vowels as well as between segments of different lengths, we restricted our analyses to CVCC words which were our most common segmental class (Fig. 1B). This resulted in a 9-by-9 auditory and VT perceptual mRDM for the CVCC trained words (Fig. 1E). These representational spaces are highly correlated (r = 0.94) and reflect the close representational congruence between auditory and VT vocoded stimuli.

#### Whole-Brain Searchlight RSA Analysis

RSA (Kriegeskorte and Kievit, 2013; Kriegeskorte et al., 2008) analyses were performed using the CoSMoMVPA toolbox (Oosterhof, Connolly, & Haxby, 2016), Surfing Toolbox (Oosterhof et al., 2011) and custom MATLAB code. Searchlights were constructed around each surface node by selecting the 30 closest voxels measured by geodesic distance. Within a given searchlight, the activity (t-statistic) in the voxels for each condition constituted its pattern. A cocktail-blank removal was performed on this condition-by-voxel data matrix whereby the mean pattern of activity across conditions was removed for each voxel (Walther et al., 2016). A neural dissimilarity matrix (nRDM) was then computed in each searchlight by computing the pairwise Pearson correlation distance (1-Pearson Correlation) between the patterns of all pairs of conditions. To assess whether a given region represented stimuli in a hypothesized format, the nRDM was compared to the mRDM. This was done by taking the Spearman Correlation between the vectorized lower triangles of the nRDM and mRDM. This correlation was then Fischer z-transformed to render the correlations more normally distributed (Kriegeskorte et al., 2008).

#### ROI-Based RSA Analysis

ROI-based RSA analyses were performed in the VT scans to test if, following training, VT stimuli engaged auditory speech representations in functionally defined ROIs identified in the auditory scans. To do so, we averaged the Fischer z-transformed correlations of searchlights in a given ROI for the four groups (pre/post x vocoded/token). We then fit these average ROI correlations with a linear mixed effects model in R using the Lme4 Package. For both ROIs, we fit the maximal model that included three main effects, all interaction terms, as well as a random slope and intercept. The random effects terms allowed us to model the subject-specific variability in the pre-training and the training-related change in correlation. (Glasser et al., 2016)The final model is shown below:

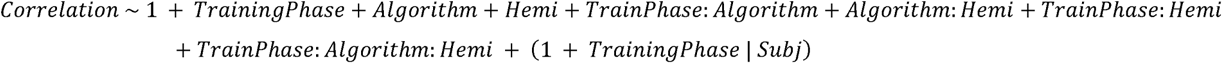

The reference group corresponding to the intercept was specified as pre-training, token-based, right-hemisphere. All βs reported reflect deviations from this reference group given the other effects. The model was estimated using REML and degrees of freedom were adjusted using the Satterthwaite approximations. Post-hoc contrasts were computed using the *emmeans* package and all reported p-values were corrected for multiple comparisons using Tukey’s method.

### Task-Related Functional Connectivity

Functional connectivity analyses were performed using the CONN-fMRI toolbox (Whitfield-Gabrieli & Nieto-Castanon, 2012). To do so, native-space functional data were smoothed using an 8mm FWHM smoothing kernel. Next, anatomical scans were segmented to identify regions of white matter and CSF. We then regressed out the signals from these regions using CompCor (Behzadi, Restom, Liau, & Liu, 2007) as well as the main effect of task. Whole-brain seed-to-voxel correlation maps were then computed within each subject. Finally, we mapped each subject’s correlation maps to a standard cortical mesh using 3dVol2Surf in order to perform group analyses.

### Whole-Brain Statistical Correction

We tested the group-level significance of whole-brain RSA analyses as well as functional connectivity differences by first computing a t-statistic at each node on the standard surface. To correct these t-statistic maps for multiple comparisons, we first estimated the smoothness of the data for each analysis in each hemisphere using the AFNI/SUMA *SURFFWHM* command. We then used this smoothness estimate to generate noise surface maps using the AFNI/SUMA *slow_surf_clustsim.py* command. This then allowed us to generate an expected cluster size distribution at various thresholds that we compared clusters in our actual data to. For the auditory scan, we performed a one-sample t-test against 0 and applied a two-tailed cluster-defining threshold of α = .001. For the functional connectivity analyses in the VT scan, we performed a two-sample paired t-test to seed-to-voxel functional connectivity in subjects pre- and post-training. We applied a two-tailed cluster-defining threshold of α = .005. All resulting clusters were corrected at the p ≤ .05 level. Tables report the coordinates of the center of mass of clusters in MNI space and their location as defined by the Glasser Atlas (Glasser et al., 2016).

## Results

### Behavior

Subjects (n=20) were trained to recognize stimuli derived from either a token-based of vocoded auditory-to-VT sensory substitution algorithm (Fig. 1C), Subjects completed 6 behavioral training sessions in which they performed a N-AFC task on each level (see Material and Methods). Only a single session was performed per day. To progress to the next level, subjects had to achieve at least 80% accuracy on the current level. Both vocoded and token-based achieved progressively higher levels in the behavioral training paradigm across training sessions (Fig. 2A). The median final levels achieved were 8 and 7 for the token-based and vocoded VT groups respectively. After the final post-training fMRI scan, subjects completed a 10-AFC test on the trained words (Fig. 2B). All subjects performed better than chance (10%) and the median accuracies were 35.3% and 48.5% for the token-based and vocoded VT groups respectively. A two-sample t-test revealed no significant difference in accuracy between algorithm groups (*t*(18) = 0.386, p = 0.704).

**Figure 2:**
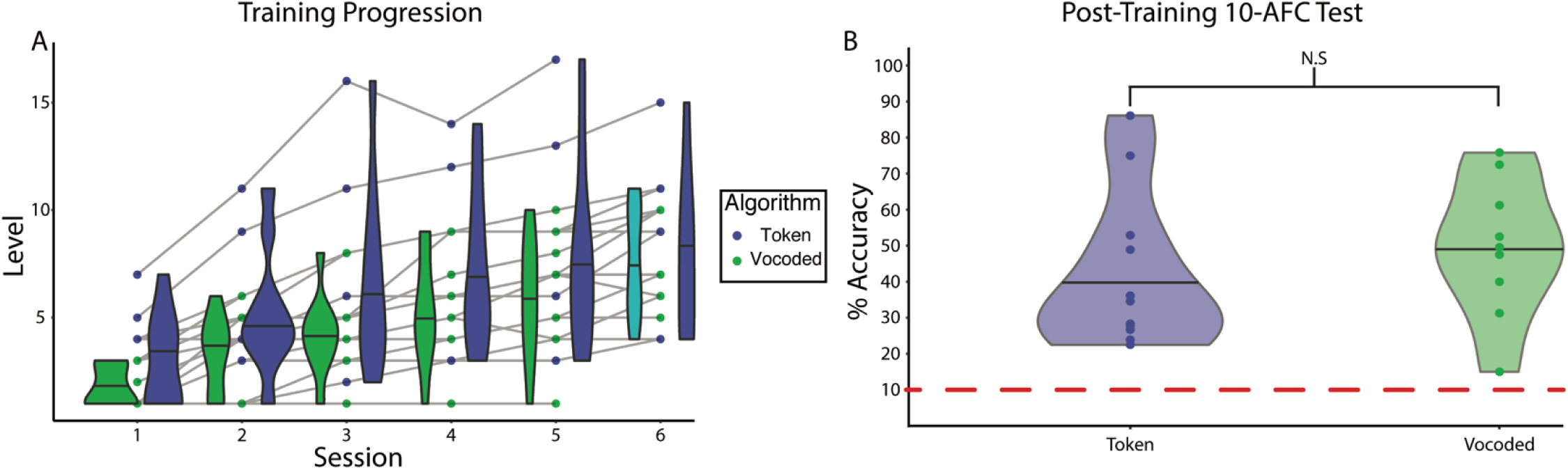
Progression of learning VT stimuli as speech. (A) Shows the leveling up of individuals on the behavioral training paradigm across sessions. Shaded lines connect the same individual across sessions. Data for the final session of two subjects was lost due to technical error. (B) Shows the performance of subjects by algorithm group on 10-AFC task completed after the final post-training fMRI scan. A two-sample t-test reveals no significant difference in performance between the groups (*t*(18) = 0.386, p = 0.704). Dashed red line indicates chance performance. Horizontal lines in the violin plots reflect the median.

### Univariate fMRI Analysis

Univariate analyses were conducted to examine the activation in response to the auditory and VT stimuli. In the auditory scan, the contrast of “All Words>Baseline” revealed bilateral Superior Temporal Gyrus (STG) activation (Table S1 and Fig. 3A). In the VT scans, unpaired two-sample t-tests revealed no significant differences between the vocoded and token-based groups in either the pre-training or post-training phase. Therefore, subjects were combined within training-phase to test for the cortical common response to VT stimulation. The contrast “All Vibrotactile Words>Baseline” revealed several regions, including bilateral supplementary motor area (SMA), precentral gyri (Table S1 and Fig. 3B-C). No significant clusters were identified for the post- vs pre- training contrast. To gain a better picture of the neuronal selectivity underlying these responses, we performed a series of RSA analyses.

**Figure 3:**
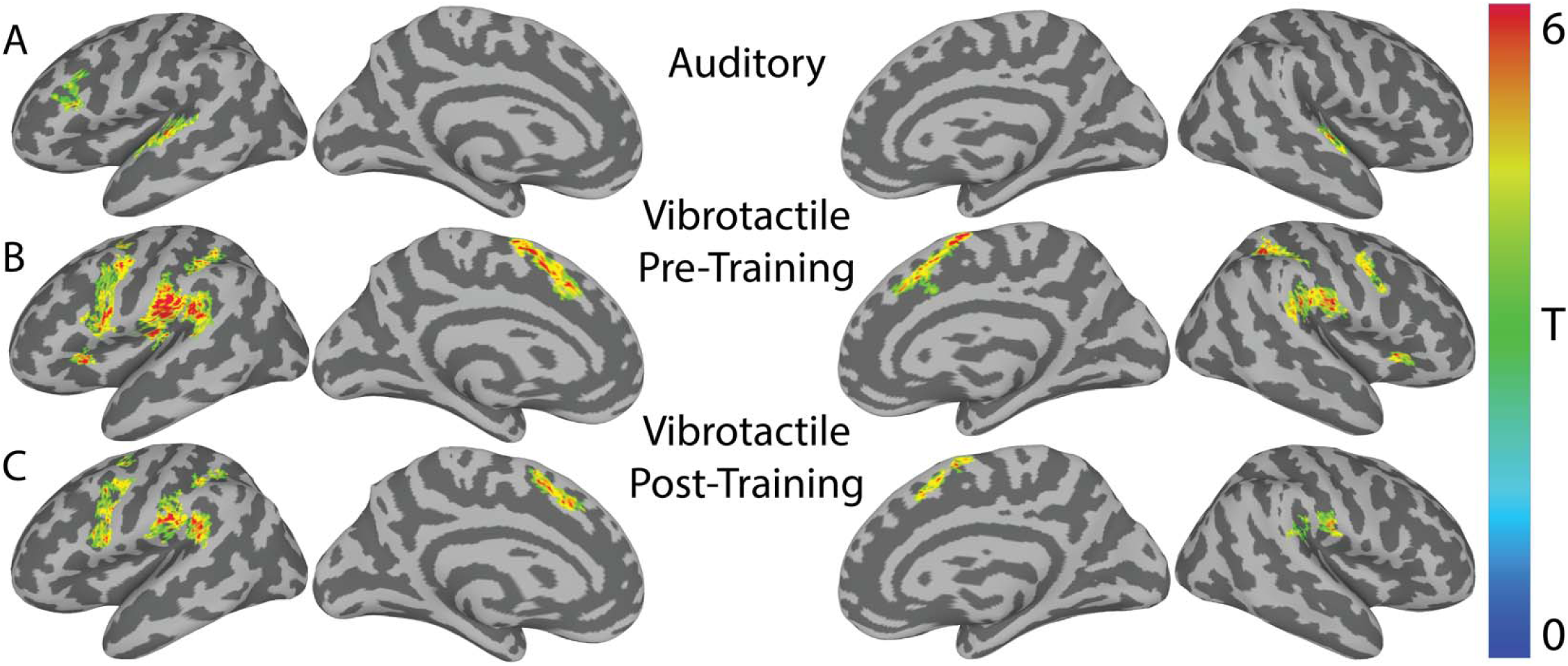
Univariate activity for “Stimuli-Baseline” in the auditory and VT scans. (A) Shows the group-level speech perception network revealed by the contrast of all auditory words > baseline. (B) Shows the pre-training group-level VT perception network revealed by the contrast of all vibrotactile words > baseline. (C) Same as (B), but for post-training scans. Results are rendered on a SUMA-derived standard surface. All results are presented at a cluster-defining two-tailed α = 0.005 and p ≤ 0.05.

### Whole-brain searchlight analysis reveals bilateral STG regions are engaged in the perception of spoken vocoded words

We conducted a whole-brain searchlight RSA analysis to identify regions showing selectivity for auditory vocoded words. In each searchlight we constructed a neural RDM that was correlated to the auditory perceptual mRDM (see Methods). The group-level t-statistic map was thresholded at a two-tailed α = .001 and the resulting clusters were corrected at two-tailed p ≤ 0.05 (Fig. 4). This revealed left (x = -58, y = -18, z = 5; α = 0.001; p = 0.001) and right mid-STG (x = 58, y = -14, z = 3; α = 0.001; p = 0.016) clusters. Of the 75 nodes in the left mid-STG cluster, 8 are in left A1, 21 are in the lateral belt, 28 are in the parabelt, and 31 are in A4 as defined by the Glasser Atlas. Of the 44 nodes in the right mid-STG cluster, 0 are in right A1, 12 are in the lateral belt, 25 are in the parabelt, and 16 are in A4. Thus, the regions identified in this analysis are non-primary auditory cortical regions that are likely selective for complex auditory spectrotemporal patterns involved in speech perception (Hamilton et al., 2020).

**Figure 4:**
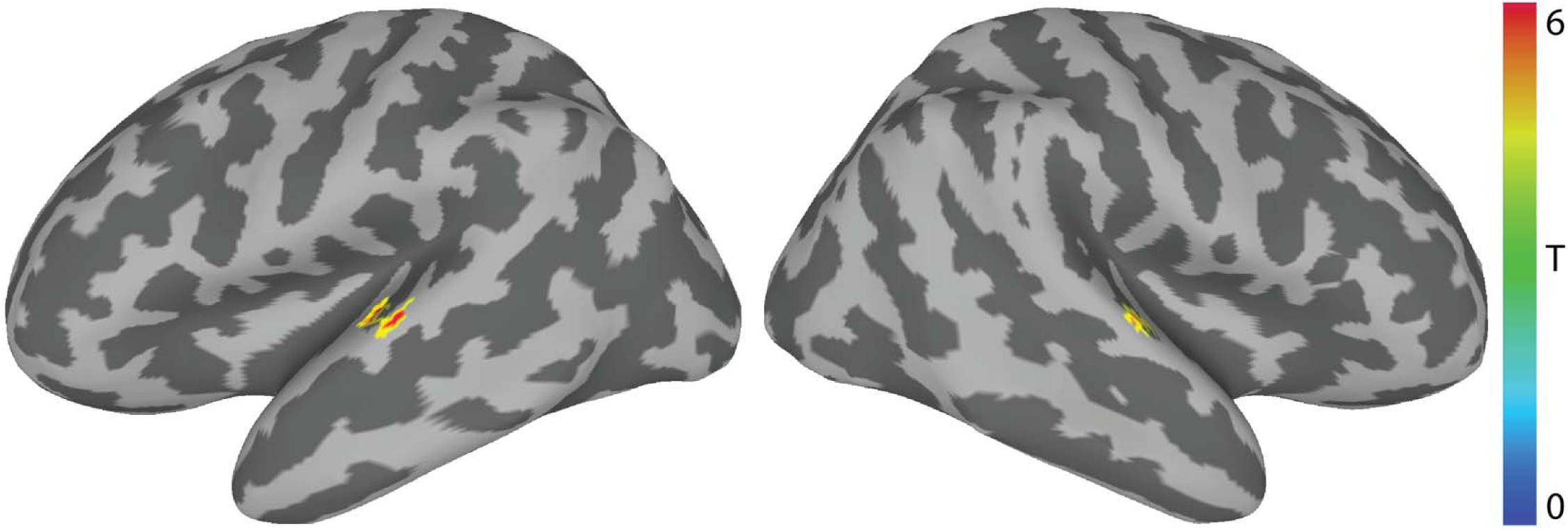
Auditory Scan – Representational similarity analysis (RSA) of vocoded auditory words. RSA revealed that neural RDMs in bilateral STG regions significantly correlated with the predicted auditory perceptual mRDM (Fig. 1E) (n=18; α = 0.001; p ≤ .05). The center of mass of the left STG cluster was centered on MNI: -58, -18, 5. The center of mass of the right STG cluster was centered on MNI: 58, -14, 3. Colors reflect across-subject t-statistics.

### ROI-based analysis reveals that the right auditory word-selective region shows selectivity for VT vocoded, but not token-based words following VT speech training

Next, we conducted ROI-based RSA analyses to test the prediction that trained VT stimuli would engage the same representations as auditory words in the mid-STG. To do so, we first computed the average Fisher transformed correlation between the vibrotactile nRDMs and the auditory perceptual mRDM for the 9 trained CVCC words in the VT scans. A linear mixed-effects model was then constructed (see Methods) to test the effects of training phase, algorithm, hemisphere, as well as the interaction among them. This analysis revealed a significant interaction effect between training phase and algorithm (β = 0.240, *t(31.09)* = 2.679, p = 0.012; Table S2). Post-hoc tests revealed a significant (*t*(31.1) = 3.380, p = 0.010 Tukey-adjusted) increase between the pre- and post-training correlations in the right mSTG for the vocoded but not (*t*(31.1) = -0.408, p = 0.977 Tukey-adjusted) the token-based group. Furthermore, post-hoc tests did not reveal a significant increase between the pre- and post-training correlations in the left mSTG for either the vocoded (*t*(31.1) = 1.781, p = 0.302 Tukey-adjusted) or the token-based (*t*(31.1) = 0.250, p = 0.994 Tukey-adjusted) group. Although there was no significant three-way interaction, we performed exploratory analyses to compare the correlation between the left vs. right mid-STG. This revealed significantly (*t*(9) = 2.783, p = 0.021) higher correlations post-training in the right than left mSTG. In addition, there was a marginally significant (*t*(9) = 2.185, p = 0.057) difference when the difference between pre- and post-training correlations were compared between the right and left mid-STG. These results indicate that trained VT stimuli based on vocoded speech engaged auditory speech representations in the mid-STG and did so more strongly than token-based VT stimuli, and there was no evidence that token-based VT stimuli engaged these auditory speech representations. Furthermore, there is evidence that this effect may be stronger in the right hemisphere than the left.

The noteworthy difference in the engagement of mid-STG auditory speech representations for the vocoded but not token-based VT stimuli raised the question what other brain areas might underlie subjects’ ability to learn the token-based VT stimuli as words (see Fig. 2). A possible explanation of the results is that because the token-based representation is not well matched to auditory speech representations (e.g., in its temporal dynamics), to learn the association between the two, the brain must rely on alternate strategies such as those used to learn arbitrary associations between pairs of stimuli. A key region involved in learning such associations is the hippocampus (McClelland et al., 1995; O’Reilly and Rudy, 2001). Therefore, we tested whether the hippocampus encoded token-based stimuli after training.

### ROI-based analysis reveals that the Left Hippocampus is engaged during perception of VT token-based, but not vocoded stimuli

We therefore next tested the hypothesis that VT speech perception training led to an encoding of the VT stimuli in the hippocampus. If trained VT speech stimuli were stored in a representation that reflected the associated auditory speech stimuli, then we would expect neural activation pattern similarity for the VT stimuli to correlate with the perceptual similarity of the auditory speech stimuli post- but not pre-training. To test this hypothesis, we correlated neural activation patterns in response to VT speech stimuli in the two different encoding schemes with the auditory perceptual mRDM before and after training. These correlations were then fit with a linear mixed effects model. This analysis revealed a significant two-way interaction between training phase and hemisphere (β = 0.095, *t(36)* = 2.696, p = 0.011; Fig. 6; Table S3) as well as a significant three-way interaction effect between training phase, algorithm, and hemisphere (β = -0.151, *t(36)* = -3.027, p = 0.005; Table S3).

**Figure 5:**
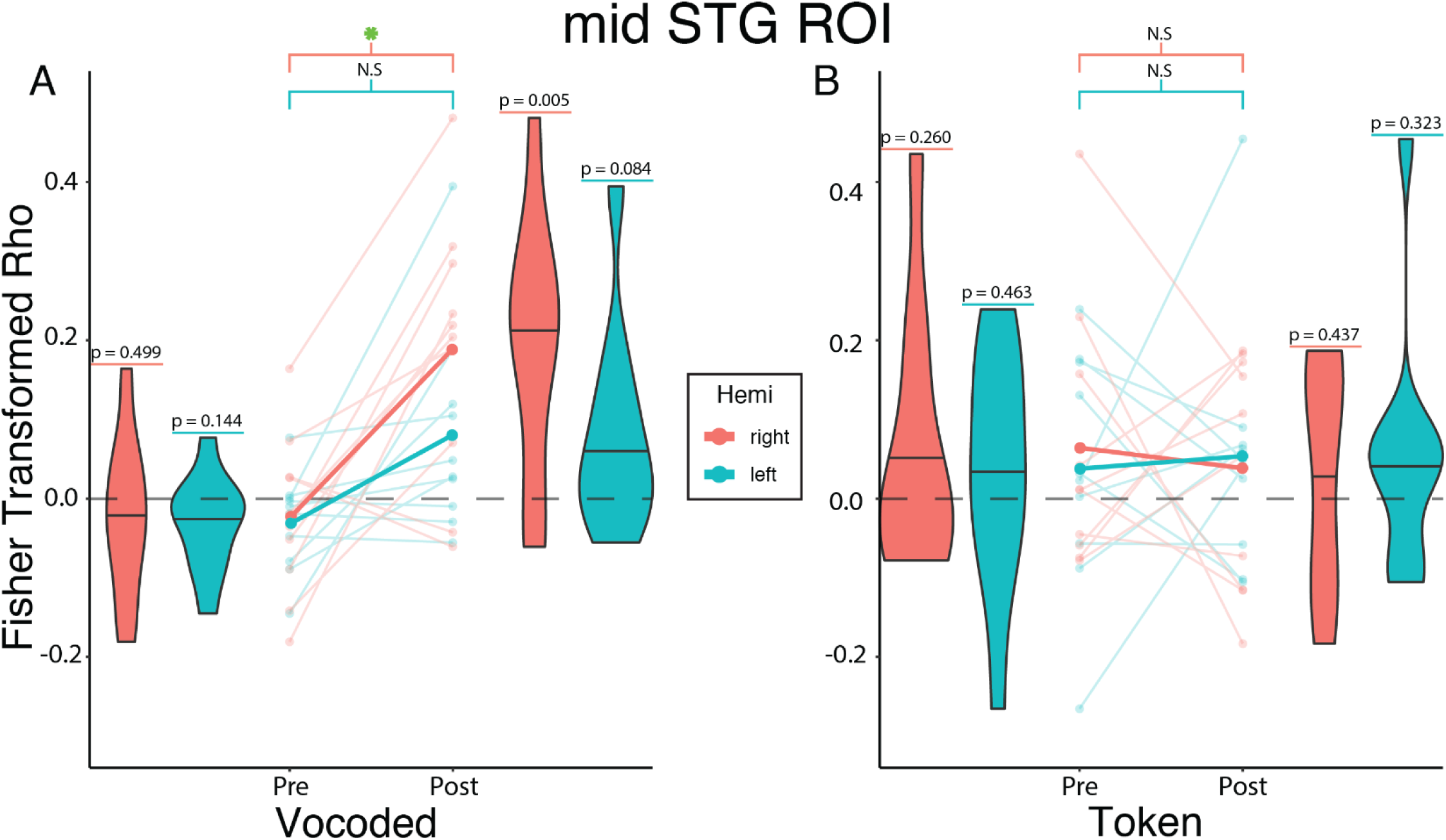
Vocoded but not token-based VT stimuli are represented in mid-STG auditory speech region following VT speech training. Linear mixed-effects analysis revealed a significant two-way interaction between Training Phase and Algorithm (β = 0.240, *t(31.1)* = 2.679, p = 0.012). To investigate this interaction, we created interaction effects plots. (A) The mean Fisher-transformed Pearson correlation between neural and model RDMs estimated from the mixed-effects model for the vocoded group are represented by the opaque lines. For the VT vocoded group, post-hoc tests show a significant difference between pre- and post-training in the right (*t*(31.1) = 3.380, p = 0.010 Tukey-adjusted) but not the left STG (*t*(31.1) = 1.781, p = 0.302 Tukey-adjusted). (B) The same as (A) but for the token-based group. Post-hoc tests show no significant difference in the right (*t*(31.1) = 0.408, p = 0.977 Tukey-adjusted) or left STG (*t*(31.1) = 0.250, p = 0.994 Tukey-adjusted). Values above each violin reflect the *uncorrected* p-value from a one-sample t-test against 0. Semi-transparent lines reflect raw individual subject correlations from either the left (teal) or right (orange) STG. Horizontal lines in the violin plots reflect the median. Green asterisk marks significant (p≤.05) differences after multiple comparisons correction.

**Figure 6:**
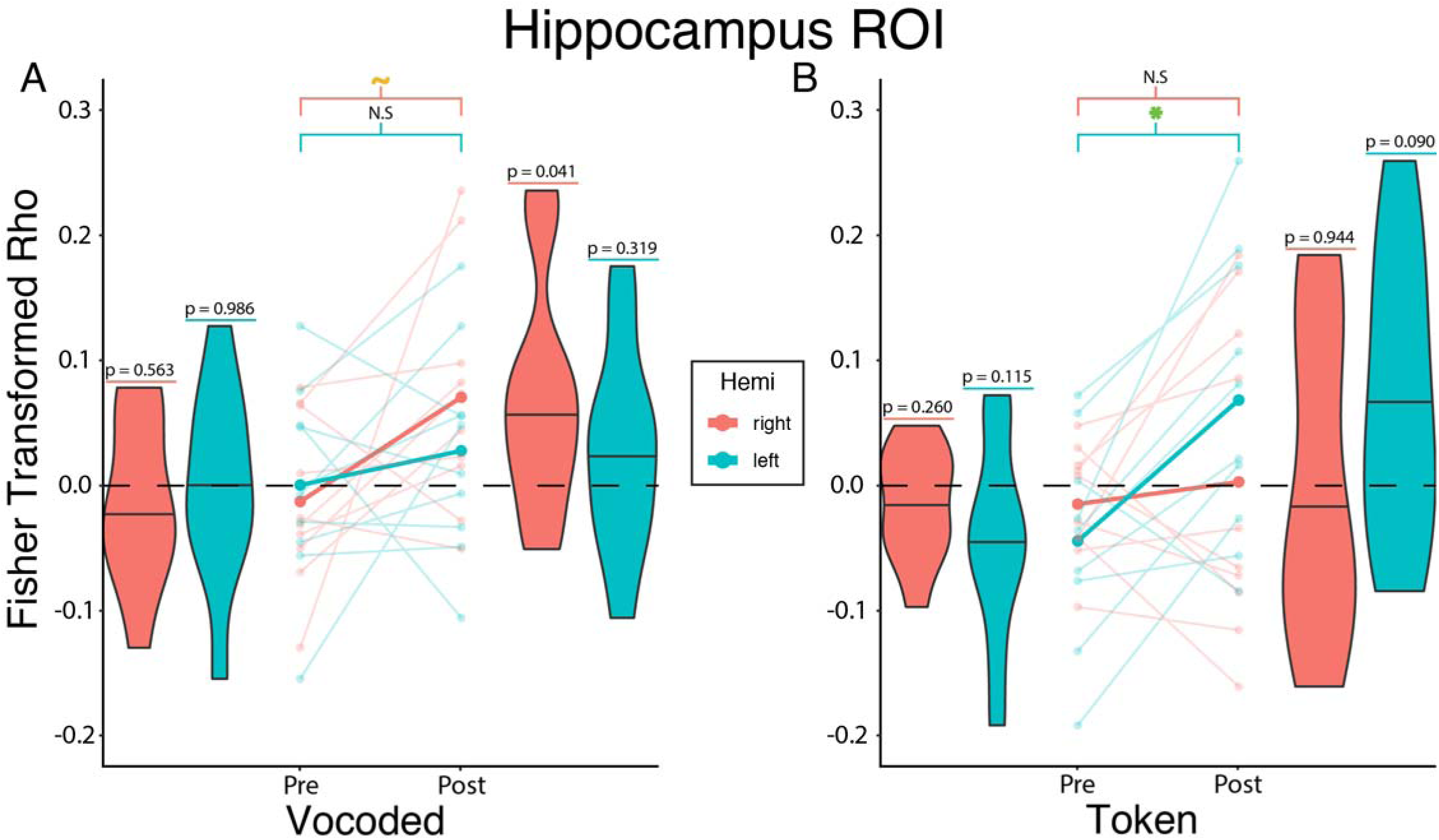
Token-based but not vocoded VT speech stimuli are represented in the left hippocampus following training. Linear mixed-effects analysis revealed a significant three-way interaction between Training Phase, Algorithm, and Hemisphere (β = -0.151, *t(36)* = -3.027, p = 0.005). To investigate this interaction, we created interaction effects plots. (A) The mean Fisher-transformed Pearson correlation between neural and model RDMs estimated from the mixed-effects model for the vocoded group are represented by the opaque lines. For the VT vocoded group, post-hoc tests show a trending difference between pre- and post-training in the right (*t*(30.7) = 2.387, p = 0.101 Tukey-adjusted) but not the left STG (*t*(30.7) = 0.785, p = 0.861 Tukey-adjusted). (B) The same as (A) but for the token-based group. Post-hoc tests show no significant difference in the right (*t*(30.7) = 0.506, p = 0.957 Tukey-adjusted), but do show a significant difference in the left STG (*t*(30.7) = 3.232, p = 0.015 Tukey-adjusted). Values above each violin reflect the uncorrected p-value from a one-sample t-test against 0. Semi-transparent lines reflect raw individual subject correlations from either the left (teal) or right (orange) STG. Horizontal lines in the violin plots reflect the median. Green asterisk and orange tilde mark significant (p≤.05) and trending (p≤.1) differences, respectively, after multiple comparisons correction.

The three-way interaction suggests that the relationship between training phase and hemisphere varied depending on the algorithm. Post-hoc tests revealed a significant (*t*(30.7) = 3.232, p = 0.0148 Tukey-adjusted) training-related increase in correlations for the token-based but not vocoded (*t*(30.7) = 0.785, p = 0.861 Tukey Adjusted) VT group in the left hemisphere. In the right hemisphere, there was a trending increase in correlation for the vocoded group (*t*(30.7) = 2.387, p = 0.101 Tukey Adjusted) but not the token-based (*t*(30.7) = .506, p = 0.957 Tukey Adjusted) VT group.

### Training with Vocoded VT Speech Stimuli Increases Functional Connectivity Between Somatosensory and Auditory Regions

Previous studies showed that learning is accompanied by increased functional connectivity between cortical areas (Lewis et al., 2009; Siuda-Krzywicka et al., 2016; Urner et al., 2013). Therefore, we tested the hypothesis that training on the vocoded VT word stimuli was associated with increased functional connectivity of somatosensory regions and the auditory word-selective right mid-STG ROI (Fig. 4). To do so, we computed the training-related changes in the right mid-STG seed-to-voxel functional connectivity in the vocoded group (Fig. 7A, Table S4). This revealed two clusters, one in the left STG (x = -50, y = -19, z = 7; α = 0.005; p = 0.044) and another in the left secondary somatosensory (SII) (x = -55, y = -28, z = 21; α = 0.005; p = 0.026). Furthermore, reasoning that VT stimulation on the right arm would engage the left SII region, we performed an additional seed-to-voxel analysis using the left SII seed defined by the Glasser atlas (Glasser et al., 2016). This complementary analysis revealed two clusters, one in the right insula and Heschl’s Gyrus (x = 40, y = -17, z = 11; α = 0.005; p = 0.001) and another in the right STG (x = 63, y = -22, z = 7; α = 0.005; p = 0.001). The left SII also showed an increase in connectivity to the left central sulcus (x = -40, y = -19, z = 42; α = 0.005; p = 0.001). Using the left mid-STG region as a seed revealed significantly increased connectivity with the right STG while using the right SII revealed significant training-related changes confined to bilateral SII. (Fig. S2, Table S4). Similar seed-to-voxel analyses also using the left hippocampus or the bilateral mid-STG ROIs as seeds revealed no significant training-related differences in the token-based group. This pattern of training-related functional connectivity between somatosensory and auditory areas for VT vocoded but not token based stimuli was also found when calculating ROI-to-ROI functional connectivity (Fig. S3). These results support a model in which vocoded VT speech training leads to increased functional connectivity between somatosensory areas and auditory speech areas.

**Figure 7:**
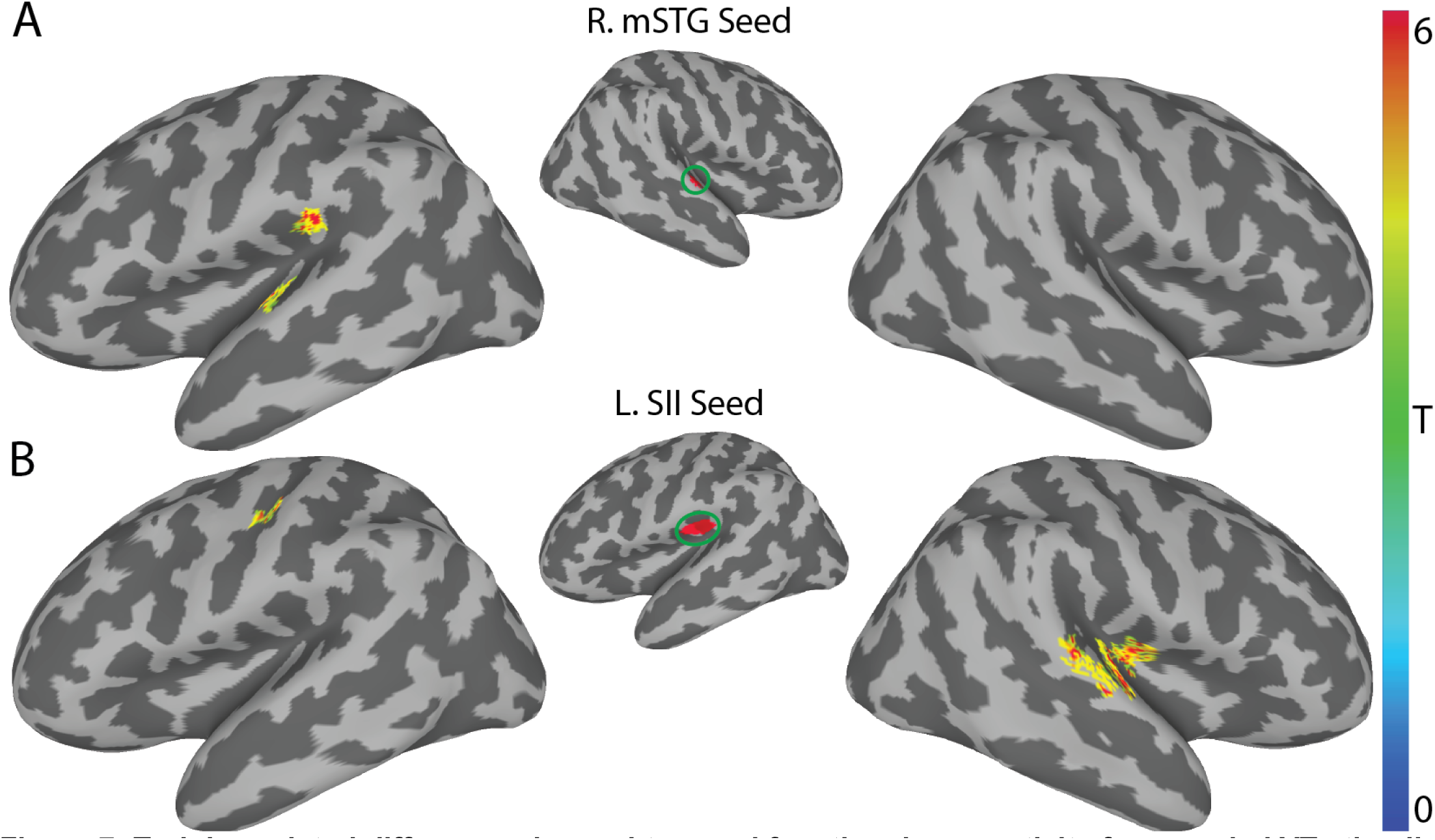
Training related differences in seed-to-voxel functional connectivity for vocoded VT stimuli. (A) Using the right mid-STG ROI (Fig. 4) as a seed revealed two significant clusters of increased functional connectivity after training in the left STG (MNI: -50, -19, 7) and in the left supramarginal gyrus (MNI: -55, -28, 21). (B) Using the left SII seed derived from the Glasser atlas revealed a significant cluster in the left central sulcus (MNI: -40, -19, 42). It also identified two significant clusters in the right hemisphere. The first encompassed right insula and Heschl’s gyrus (MNI: 40, -17, 11). The other is on the right STG (MNI: 63, -22, 7). All results shown are corrected at two-tailed voxel-wise α = 0.005 and cluster-p ≤ 0.05. Colors reflect across-subject t-statistics.

## Discussion

Metamodal theories of brain organization (Heimler et al., 2015; Pascual-Leone and Hamilton, 2001) propose that cortical areas are best described by their task-specific sensory modality-invariant function. However, mixed evidence for metamodal brain organization in neurotypical individuals (Amedi et al., 2007; Bola et al., 2017; Ptito et al., 2005; Sadato et al., 1996; Siuda-Krzywicka et al., 2016) has raised the question of if and under what conditions metamodal engagement occurs. We argue, based on theoretical considerations, that testing the metamodal hypothesis requires not just a consideration of high-level tasks (Marr’s (Marr, 1982) top level of “computational theory”) but also and critically their algorithmic implementation (Marr’s second level). In the current study, we investigated this hypothesis by training subjects on the same task (recognition of vibrotactile stimuli derived from auditory words) using one of two different auditory-to-VT sensory substitution algorithms. One algorithm (vocoded) preserved the temporal modulations of auditory speech while the other algorithm (token) attempted to establish an abstract congruence between VT patterns and the phonetic features found in speech. First, using whole-brain searchlight RSA we identified auditory perceptual speech representations whose locations along the superior temporal gyrus are compatible with models of the auditory ventral speech recognition stream (DeWitt and Rauschecker, 2012; Hickok and Poeppel, 2007; Rauschecker and Scott, 2009). Notably, this speech selectivity was found bilaterally, in agreement with other models of speech processing in the brain (Hickok and Poeppel, 2007). We then showed that, before training, neither the vocoded nor the token-based VT stimuli selectively engaged these auditory speech areas, as expected. Next, over the course of six behavioral sessions, we trained two groups of subjects to recognize the VT-encoded word stimuli, with each group trained on a different encoding scheme. Both groups of subjects achieved comparable levels of proficiency, eliminating performance differences as a reason for the different training effects at the neural level. Crucially, RSA revealed that after training, only the vocoded but not the token-based VT stimuli engaged an auditory-speech selective region in the mid-STG (Hamilton et al., 2020). In addition, both encoding schemes (to different degrees) appeared to engage hippocampal areas previously implicated in paired-associate learning. Finally, we found evidence that metamodal engagement of the mid-STG by vocoded VT stimuli was associated with a training-related increase in functional coupling between the mid-STG and secondary somatosensory areas. Evidence of training-related increases in functional coupling was not found for token-based stimuli.

In this study, we show that adequately capturing (and eventually harnessing) the metamodal potential of cortex requires not only the right task and sensory modalities but also an understanding of the information representation in these regions. Prior work has primarily investigated metamodal engagement in congenitally sensory-deprived individuals (Arno et al., 2001; Bola et al., 2017; Lomber et al., 2010; Ptito et al., 2005; Reich et al., 2011; Sadato et al., 1996). In such cortical areas, given the right task-relevant connectivity, bottom-up input from another sensory modality can conceivably drive the *de novo* learning of task-relevant representations even for encoding schemes very different from those in neurotypical individuals (Striem-Amit et al., 2012). However, in neurotypical adults, existing representations in traditionally unisensory areas reflect the task-relevant features of the typical sensory input (Lewicki, 2002; Simoncelli and Olshausen, 2001). Therefore, for metamodal engagement to occur, information partially processed in one sensory hierarchy needs to interface with pre-existing representations derived from the typical modality. The lack of evidence for metamodal engagement of the mid-STG by token-based VT stimuli in our study and the mixed evidence in prior studies of neurotypical individuals may reflect a failure to perform this interface mapping.

The ability to map between representational formats in different sensory hierarchies likely depends on both anatomical and functional convergence. Anatomical tracer (Mothe et al., 2006a; Schroeder et al., 2003; Smiley et al., 2007) and studies in non-human primates (Kayser et al., 2009; Schroeder et al., 2001) as well as neuroimaging studies in humans (Foxe et al., 2002; Ro et al., 2013) have established convergence points between somatosensory and auditory cortices including belt and parabelt areas. Given this connectivity, prior computational studies have shown that the mapping between different representational formats can be learnt through simple biologically plausible learning rules (Davison and Frégnac, 2006; Pouget and Sejnowski, 1997; Pouget and Snyder, 2000). Still, while it is simple to learn the mapping between static features, it is non-trivial to match the temporal dynamics between functional hierarchies. For example, Davison and Frégnac (2006) computationally demonstrated the importance of temporally coherent activity between representational formats when learning the mapping between cross-modal temporal sequences using spike-timing-dependent plasticity mechanisms. In the auditory cortex specifically, studies (Moore and Woolley, 2019; Overath et al., 2015) have shown that auditory stimuli that do not preserve the same temporal modulations found in conspecific communication signals (e.g., speech, birdsong, etc.) sub-optimally drive higher-order auditory cortex and preclude learning. Recent human intracranial EEG studies (Hamilton et al., 2018; Hullett et al., 2016) have demonstrated that middle superior temporal cortex is characterized by very short temporal receptive fields necessitating relatively rapid changes in the somatosensory signal. Accordingly, we find, in the current study, that only vocoded stimuli that preserve these fast temporal dynamics are able to drive auditory perceptual speech representations in the mid-STG. Conversely, the different dynamics (see Materials and Methods) of token-based VT stimuli relative to auditory speech may explain why these stimuli were unable to interface with mid-STG speech representations.

Intriguingly, we find stronger evidence of metamodal engagement by VT vocoded stimuli in the right rather than left mid-STG. A significant body of work (Albouy et al., 2020; Boemio et al., 2005; Flinker et al., 2019; Giraud and Poeppel, 2012; Obleser et al., 2008; Zatorre and Belin, 2001) suggests that the left and right STG are differentially sensitive to spectrotemporal content of auditory stimuli. Specifically, it has been proposed (Flinker et al., 2019) that the left STG tends to sample auditory information on fast and slow timescales while the right preferentially does the latter. In the current study, our VT vocoded stimuli preserve the coarse temporal dynamics of auditory speech, but due to hardware limitations have a lower temporal resolution than the auditory source signal. In addition, the temporal resolution of vibrotactile perception is lower than that of auditory processing, with receptors in the skin acting as an additional low pass filter (Bensmaïa and Hollins, 2005). Thus, the observed metamodal coupling with the right rather than the left STG provides intriguing support for the asymmetric spectrotemporal modulation theory of hemispheric processing (Flinker et al., 2019).

Given that subjects were able to learn token-based and vocoded VT stimuli as words with roughly equal proficiency, how do token-based stimuli engage spoken word representations? Due to the slower temporal dynamics of token-based stimuli, we initially hypothesized that these stimuli may map onto higher order speech representations in areas such as the superior temporal sulcus (STS) or anterior STG that integrate temporal information on longer timescales (Hullett et al., 2016; Overath et al., 2015). However, we did not find evidence for this in the current study. An anatomical tracer study by De La Mothe (Mothe et al., 2006b) showed strong evidence of connectivity between somatosensory cortex and mid and posterior but not anterior superior temporal areas. Thus, a homologous lack of connectivity between somatosensory and anterior superior temporal areas in humans may explain why we observed no engagement of those areas after training. However, we did find evidence that token-based stimuli engage neural representations in the left hippocampus. This result fits with previous proposals that learned associations can be retrieved using paired-associate recall circuits in the medial temporal lobe (Miyashita, 2019). A more thorough understanding of this process through future studies will shed additional insight into which pathways and mechanisms are leveraged to learn different types of associations.

Previous studies have suggested that metamodal engagement is a result of top-down processes such as mental imagery rather than bottom-up processes (Lacey et al., 2009). However, given that in our study, subjects in both algorithm groups were equally proficient at recognizing VT stimuli as words, mental-imagery accounts (Borst and Gelder, 2016; Li et al., 2020; Oh et al., 2013; Tian et al., 2018) in this case would predict that both groups should engage auditory perceptual representations in the mid-STG. Yet, we found no evidence that the token-based VT stimuli engaged this area after training in the same way as auditory speech (see also (Siuda-Krzywicka et al., 2016; Striem-Amit et al., 2012)). Thus, it is unlikely that metamodal engagement of the mid-STG by vocoded stimuli is driven by top-down mechanisms.

Most prior studies (Amedi et al., 2002, 2007; Reich et al., 2011; Siuda-Krzywicka et al., 2016; Striem-Amit et al., 2012, 2015; Vetter et al., 2020) have demonstrated metamodal engagement in visual cortex. Our study extends these findings to show that metamodal engagement is possible in auditory cortex as well. To our knowledge, metamodal engagement of auditory cortex has been limited to posterior auditory association cortex (pSTS) and has only been found in congenitally deaf but not hearing individuals (Benetti et al., 2017, 2020; Bola et al., 2017; Twomey et al., 2017). Furthermore, these studies did not find evidence of metamodal engagement in neurotypical individuals. In contrast, our study provides novel evidence for metamodal engagement of intermediate auditory areas. This is particularly noteworthy given the sparse evidence for metamodal engagement of intermediate sensory areas (Heimler and Amedi, 2020). The dearth of evidence is likely due to a lack of knowledge of the structure of stimulus representations in these regions, which our work suggests is critical for successful metamodal engagement.

In summary, our results provide further evidence for the metamodal theory and advance it by demonstrating the importance of matching representational formats between functional hierarchies for achieving metamodal engagement. In particular, our results suggest that matching the temporal dynamics of representations is an important consideration when considering the feasibility of learning the appropriate mapping. This extends theories (Heimler et al., 2015; Pascual-Leone and Hamilton, 2001) that emphasize a cross-modal congruences at the Computational/Task level by additionally highlighting the need for an algorithmic congruence. Taking this need for algorithmic congruence into account may provide insight into how the brain learns to map between various levels of different functional hierarchies like sub-lexical and lexical orthography and phonology (Share, 1999). Furthermore, it suggests that therapeutic sensory substitution devices might benefit from different designs for patients with acquired rather than congenital sensory deprivation. For the former group, careful consideration should be given to the type of sensory substitution device that best interfaces with spared sensory representations. The ability to “piggyback” onto an existing processing hierarchy (e.g., auditory speech recognition) may facilitate the rapid learning of novel stimuli presented through a spared sensory modality (e.g., VT). Here we demonstrate that an algorithm (vocoding) that improves this interfacing is able to more efficiently convey the same information than an algorithm (token) that does not. Future work should explore whether this observed integration into existing processing streams leads to improved generalization and transfer of learning.

## Acknowledgments

A portion of the funding for this research was provided by Facebook. We would also like to acknowledge Ali Israr, Frances Lau, Keith Klumb, Robert Turcott, and Freddy Abnousi for their involvement in the early stages of the project, including the design and evaluation of the token-based encoding scheme. Finally, we would like to acknowledge Dr. Ella Striem-Amit for helpful feedback on earlier versions of this manuscript.

---

**Supplementary Figure 1:**
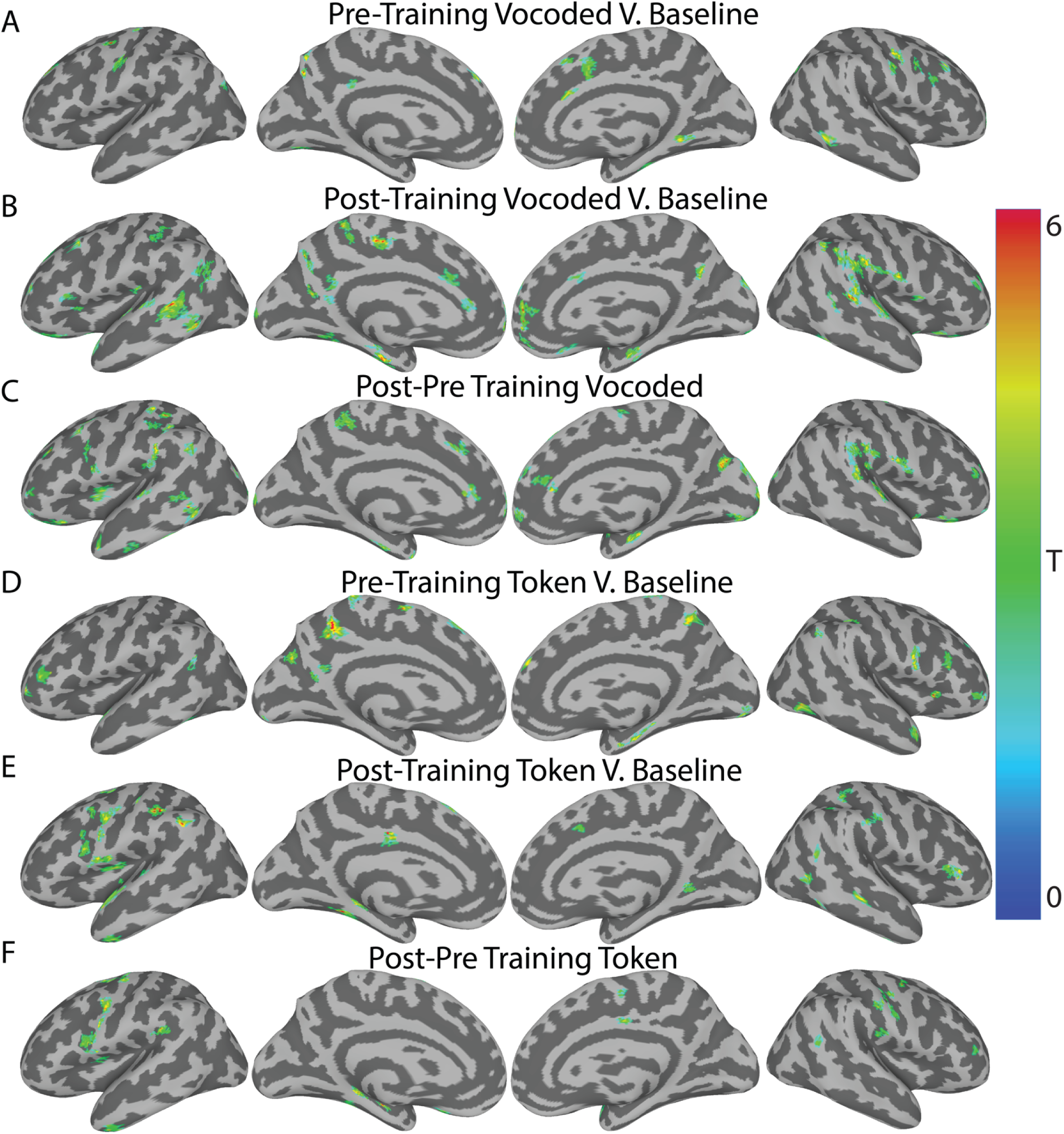
Whole-brain Searchlight RSA of Trained CVCC VT Stimuli with the Auditory Perceptual mRDM. (A-C) Whole-brain RSA results for the VT vocoded stimuli. (A) Pre-training scan: Fischer transformed correlation against 0. (B) Post-training scan: Fischer transformed correlation against 0. (C) Post minus Pre-Training change in the Fischer transformed correlations. (D-F) same as (A-C) but for the token-based stimuli. All results are at an uncorrected two-tailed voxel-wise threshold α = 0.05 with a extent threshold of 50mm^2^. Colors reflect across-subject t-statistics.

**Supplementary Figure 2:**
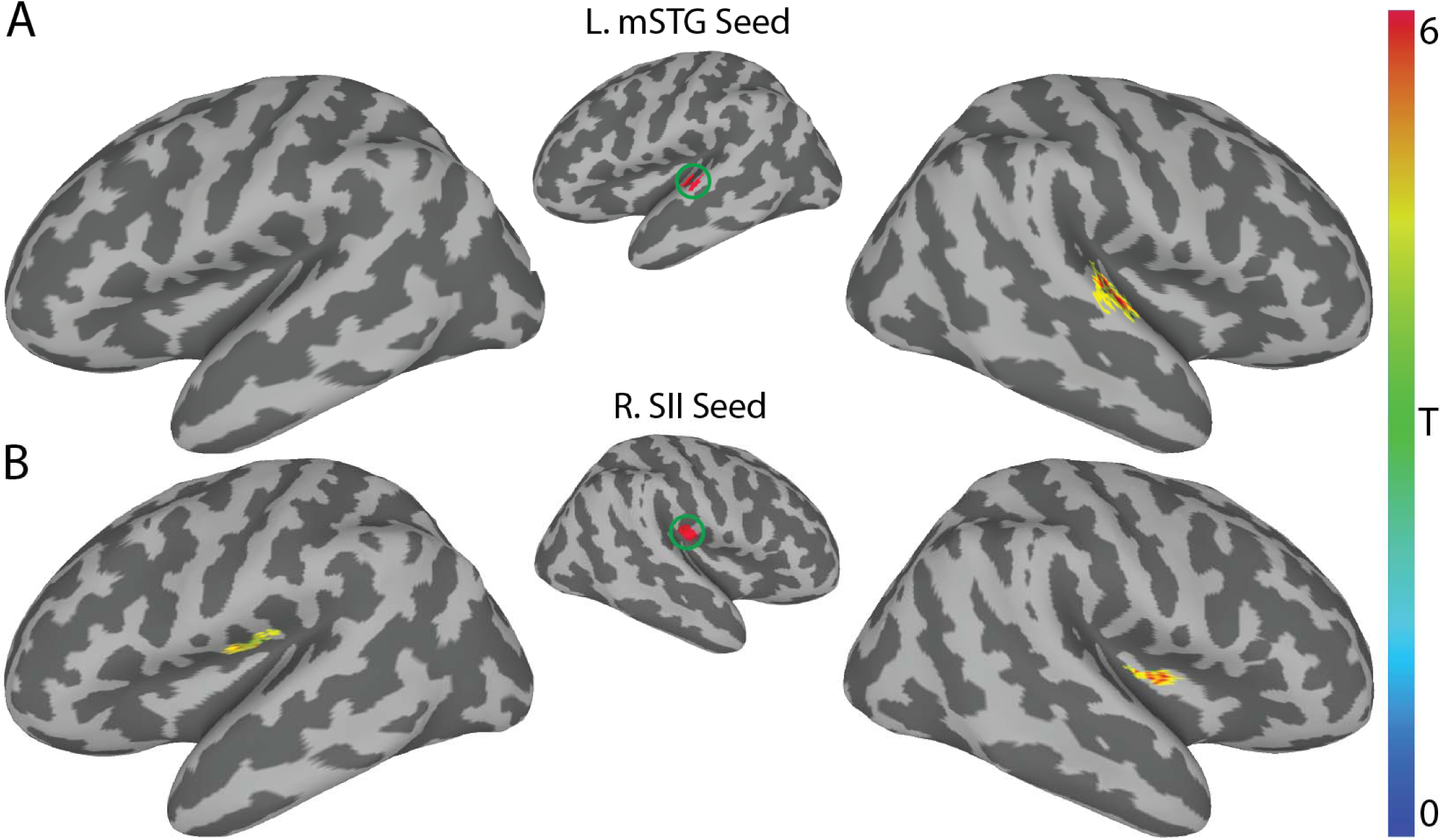
Training-related differences in seed-to-voxel functional connectivity for vocoded VT stimuli using the left STG and right SII seeds. (A) Using the left mid-STG ROI (Fig. 4) as a seed revealed one significant cluster of increased functional connectivity after training in the right mid-STG (MNI: 55, -16, 3). (B) Using the right SII seed derived from the Glasser atlas revealed a significant cluster in the left opercular region (MNI: -42, -15, 20) and right posterior Insula (MNI: 37, -3, 7). All results shown are corrected at two-tailed voxel-wise α = 0.005 and cluster-p ≤ 0.05. Colors reflect across-subject t-statistics.

**Supplementary Figure 3:**
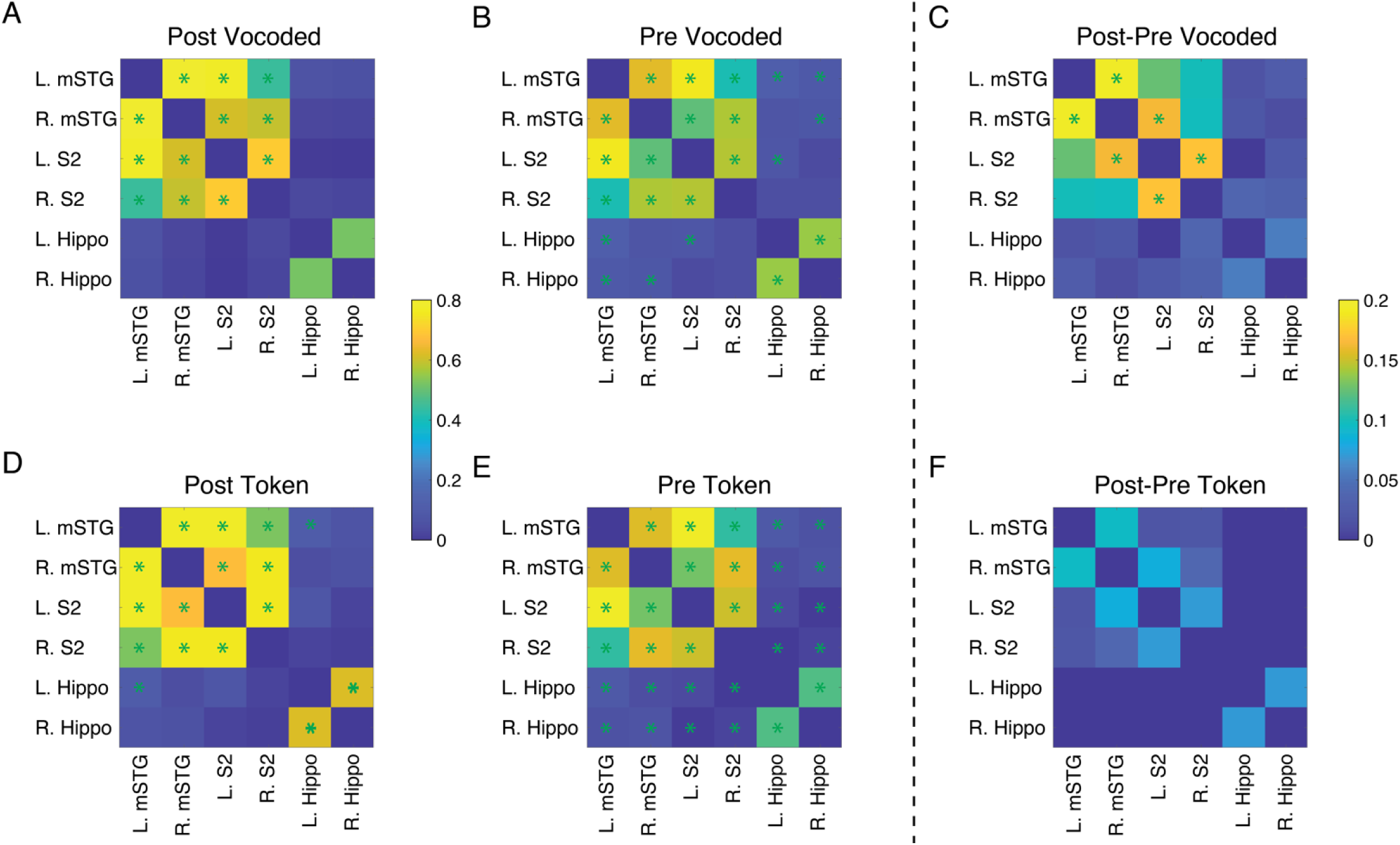
ROI-to-ROI based functional connectivity reveals significantly increased coupling between the auditory and somatosensory system after training on VT vocoded stimuli. (A-B) Shows the ROI-to-ROI functional connectivity for the VT vocoded-based group during post (A) and pre (B) training scans. (D-E) Same as (A-B) but for the VT token-based group. Color bar reflects the Fischer-transformed Pearson correlation between ROIs. A paired t-test was performed to compare changes in functional connectivity relative to baseline. Green asterisks mark p ≤ 0.05 FDR corrected. (C, F) Shows the post-pre training correlations for the VT vocoded and token-based groups respectively. Color bar reflects the Post-Pre training difference between ROIs. A paired t-test was performed to compare changes in functional connectivity post-pre training. Green asterisks mark p ≤ 0.05 FDR corrected.

**Supplementary Table 1:**
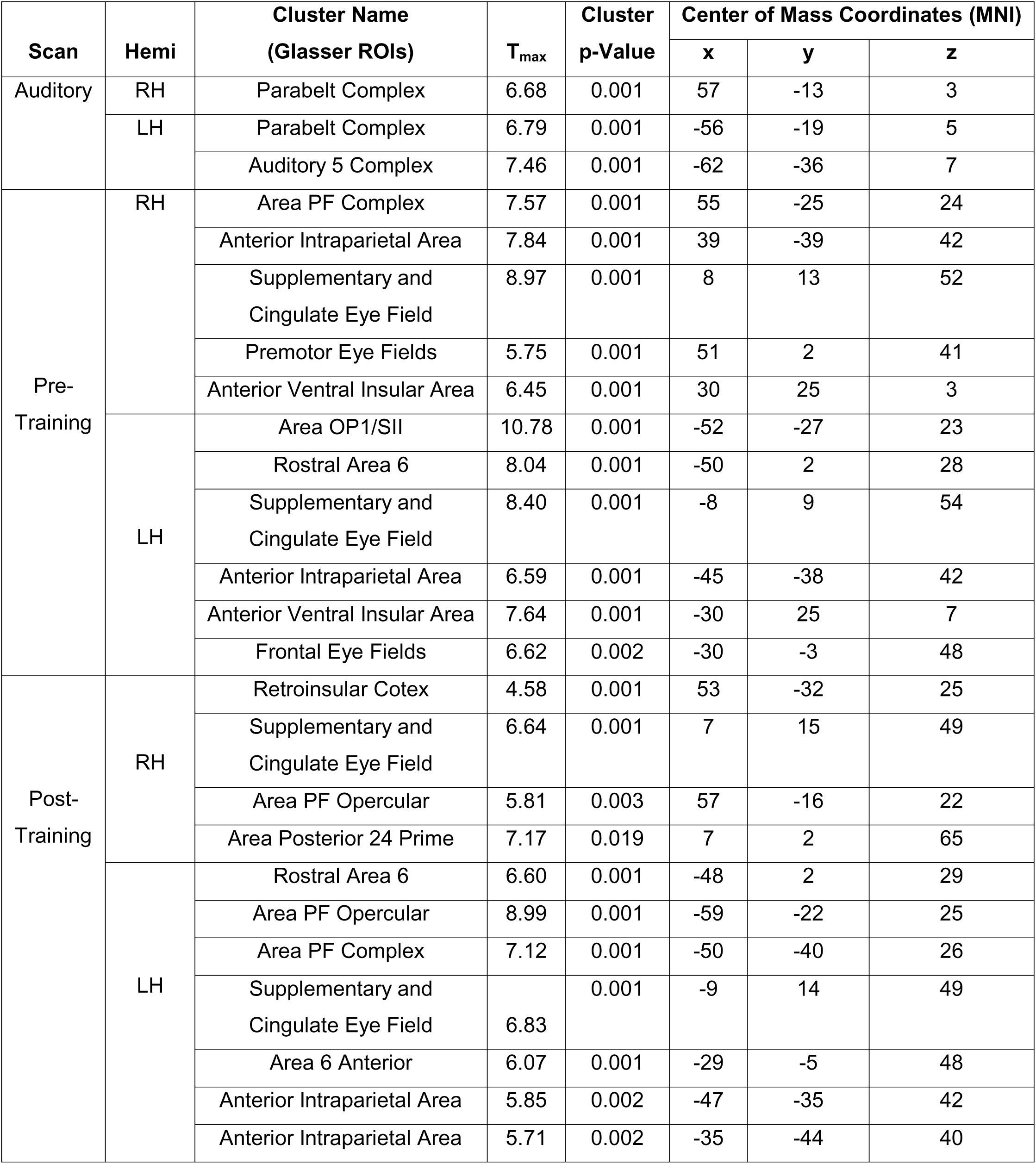
Univariate activity for all stimuli > baseline in the different scans.

| Scan | Hemi | Cluster Name<br>(Glasser ROIs) | T<sub>max</sub> | Cluster<br>p-Value | Center of Mass Coordinates (MNI) |  |  |
| --- | --- | --- | --- | --- | --- | --- | --- |
|  |  |  |  |  | x | y | z |
| Auditory | RH | Parabelt Complex | 6.68 | 0.001 | 57 | -13 | 3 |
|  | LH | Parabelt Complex | 6.79 | 0.001 | -56 | -19 | 5 |
|  |  | Auditory 5 Complex | 7.46 | 0.001 | -62 | -36 | 7 |
| Pre-Training | RH | Area PF Complex | 7.57 | 0.001 | 55 | -25 | 24 |
|  |  | Anterior Intraparietal Area | 7.84 | 0.001 | 39 | -39 | 42 |
|  |  | Supplementary and<br>Cingulate Eye Field | 8.97 | 0.001 | 8 | 13 | 52 |
|  |  | Premotor Eye Fields | 5.75 | 0.001 | 51 | 2 | 41 |
|  |  | Anterior Ventral Insular Area | 6.45 | 0.001 | 30 | 25 | 3 |
|  | LH | Area OP1/SII | 10.78 | 0.001 | -52 | -27 | 23 |
|  |  | Rostral Area 6 | 8.04 | 0.001 | -50 | 2 | 28 |
|  |  | Supplementary and<br>Cingulate Eye Field | 8.40 | 0.001 | -8 | 9 | 54 |
|  |  | Anterior Intraparietal Area | 6.59 | 0.001 | -45 | -38 | 42 |
|  |  | Anterior Ventral Insular Area | 7.64 | 0.001 | -30 | 25 | 7 |
|  |  | Frontal Eye Fields | 6.62 | 0.002 | -30 | -3 | 48 |
| Post-Training | RH | Retroinsular Cotex | 4.58 | 0.001 | 53 | -32 | 25 |
|  |  | Supplementary and<br>Cingulate Eye Field | 6.64 | 0.001 | 7 | 15 | 49 |
|  |  | Area PF Opercular | 5.81 | 0.003 | 57 | -16 | 22 |
|  |  | Area Posterior 24 Prime | 7.17 | 0.019 | 7 | 2 | 65 |
|  | LH | Rostral Area 6 | 6.60 | 0.001 | -48 | 2 | 29 |
|  |  | Area PF Opercular | 8.99 | 0.001 | -59 | -22 | 25 |
|  |  | Area PF Complex | 7.12 | 0.001 | -50 | -40 | 26 |
|  |  | Supplementary and<br>Cingulate Eye Field | 6.83 | 0.001 | -9 | 14 | 49 |
|  |  | Area 6 Anterior | 6.07 | 0.001 | -29 | -5 | 48 |
|  |  | Anterior Intraparietal Area | 5.85 | 0.002 | -47 | -35 | 42 |
|  |  | Anterior Intraparietal Area | 5.71 | 0.002 | -35 | -44 | 40 |

**Supplementary Table 2:** Linear Mixed-Effects Model Summary for the mid-STG ROIs.

| Summary of Linear Mixed Effects Model: mid-STG ROIs |  |  |  |  |  |
| --- | --- | --- | --- | --- | --- |
| Fixed Effects |  |  |  |  |  |
| Predictors | β Estimate | Confidence Interval | T-Statistic | DOF | p-value |
| Intercept | 0.063 | -0.02 – 0.14 | 1.566 | 35.34 | 0.126 |
| Training Phase | -0.026 | -0.15 – 0.10 | -0.408 | 31.09 | 0.686 |
| Algorithm | -0.086 | -0.20 – 0.03 | -1.507 | 35.34 | 0.141 |
| Hemisphere | -0.026 | -0.12 – 0.06 | -0.578 | 36 | 0.567 |
| Training Phase:Algorithm | 0.240 | 0.06 – 0.41 | 2.679 | 31.09 | 0.012 |
| Algorithm:Hemisphere | 0.018 | -0.11 – 0.14 | 0.277 | 36 | 0.783 |
| Training Phase:Hemisphere | 0.042 | -0.09 – 0.17 | 0.643 | 36 | 0.524 |
| Training Phase:<br>Algorithm:Hemisphere | -0.143 | -0.32 – 0.04 | -1.559 | 36 | 0.128 |
| Random Effects |  |  |  |  |  |
| Groups | Effect Name | σ (std. deviation) | Variance | Correlation Structure |  |
| Subj | Intercept | 0.075 | 0.006 | N/A | -0.60 |
|  | Training<br>Phase | 0.138 | 0.019 | -0.60 | N/A |
| Residual |  | 0.102 | 0.010 |  |  |

**Supplementary Table 3:** Linear Mixed-Effects Model Summary for the Hippocampus ROIs.

| Summary of Linear Mixed Effects Model: Hippocampus ROIs |  |  |  |  |  |
| --- | --- | --- | --- | --- | --- |
| Fixed Effects |  |  |  |  |  |
| Predictors | β Estimate | Confidence Interval | T-Statistic | DOF | p-value |
| Intercept | -0.015 | -0.06 – 0.03 | -0.668 | 34.94 | 0.508 |
| Training Phase | 0.018 | -0.05 – 0.09 | 0.506 | 30.67 | 0.616 |
| Algorithm | 0.002 | -0.06 – 0.07 | 0.066 | 34.94 | 0.948 |
| Hemisphere | -0.029 | -0.08 – 0.02 | -1.185 | 36 | 0.244 |
| Training Phase:Algorithm | 0.066 | -0.04 – 0.17 | 1.330 | 30.67 | 0.193 |
| Algorithm:Hemisphere | 0.043 | -0.03 – 0.11 | 1.211 | 36 | 0.234 |
| Training Phase:Hemisphere | 0.095 | 0.02 – 0.17 | 2.696 | 36 | 0.011 |
| Training Phase:<br>Algorithm:Hemisphere | -0.151 | -0.25 – -0.05 | -3.027 | 36 | 0.005 |
| Random Effects |  |  |  |  |  |
| Groups | Effect Name | σ (std. deviation) | Variance | Correlation Structure |  |
| Subj | Intercept | 0.042 | 0.002 | N/A | 0.03 |
|  | Training<br>Phase | 0.077 | 0.006 | 0.03 | N/A |
| Residual |  | 0.056 | 0.003 |  |  |

**Supplementary Table 4:** Training-related changes in functional connectivity in the vocoded group.

| Seed ROI | Hemi | Cluster Location<br>(Glasser ROIs) | T<sub>max</sub> | Cluster<br>p-Value | Center of Mass Coordinates (MNI) |  |  |
| --- | --- | --- | --- | --- | --- | --- | --- |
|  |  |  |  |  | x | y | z |
| IS2 | RH | Insular Granular Complex | 8.04 | 0.001 | 40 | -17 | 11 |
|  |  | Auditory 5 Complex | 8.44 | 0.001 | 63 | -22 | 7 |
|  | LH | Primary Motor Cortex | 7.73 | 0.012 | -40 | -19 | 42 |
| ISTG | RH | Lateral Belt Complex | 6.54 | 0.001 | 53 | -18 | 6 |
| rS2 | RH | Posterior Insular Area 2 | 7.41 | 0.017 | 37 | -8 | 6 |
|  | LH | Area OP2-3/VS | 5.73 | 0.026 | -42 | -16 | 20 |
| rSTG | LH | Area PF<sub>cm</sub> | 8.20 | 0.026 | -55 | -28 | 21 |
|  |  | Lateral Belt Complex | 6.75 | 0.044 | -50 | -19 | 7 |

